# Mutation of a PER2 phosphodegron perturbs the circadian phosphoswitch

**DOI:** 10.1101/2019.12.16.876615

**Authors:** Shusaku Masuda, Rajesh Narasimamurthy, Hikari Yoshitane, Jae Kyoung Kim, Yoshitaka Fukada, David M. Virshup

## Abstract

Casein kinase 1 (CK1) plays a central role in regulating the period of the circadian clock. In mammals, PER2 protein abundance is regulated by CK1-mediated phosphorylation and proteasomal degradation. On the other hand, recent studies have questioned whether the degradation of the core circadian machinery is a critical step in clock regulation. Prior cell-based studies found that CK1 phosphorylation of PER2 at Ser478 recruits the ubiquitin E3 ligase β-TrCP, leading to PER2 degradation. Creation of this phosphodegron is regulated by a phosphoswitch that is also implicated in temperature compensation. However, *in vivo* evidence that this phosphodegron influences circadian period is lacking. Here, we generated and analyzed PER2-Ser478Ala knock-in mice. The mice showed longer circadian period in behavioral analysis. Molecularly, mutant PER2 protein accumulated in both the nucleus and cytoplasm of the mouse liver, while *Per2* mRNA levels were minimally affected. Nuclear PER1, CRY1 and CRY2 proteins also increased, probably due to stabilization of PER2-containing complexes. In mouse embryonic fibroblasts derived from PER2-Ser478Ala::LUC mice, three-phase decay and temperature compensation of the circadian period was perturbed. These data provide direct *in vivo* evidence for the importance of phosphorylation-regulated PER2 stability in the circadian clock and validate the phosphoswitch in a mouse model.

## Introduction

The cell-autonomous clock systems oscillating with a period of ~24 h have evolved to coordinate physiological functions with daily environmental changes [1]. The molecular mechanism at the heart of the circadian clock is based on transcriptional/translational feedback loops that consist of clock genes and the encoded clock proteins. In the mammalian core clockwork, two proteins, circadian locomotor output cycles kaput (CLOCK) and brain muscle arnt-like 1 (BMAL1), form a heterodimer that binds to the cis-regulatory DNA element E-boxes to activate transcription of *Period (Per)* and *Cryptochrome (Cry)* genes. Translated PER and CRY proteins form a multimeric complex with casein kinase 1 (CK1δ/ε) in the cytoplasm, and then translocate to the nucleus, where they inhibit the transcriptional activity of CLOCK-BMAL1 complex [2–4]. The key molecular components of the clock in all organisms are subject to various posttranslational modifications, most notably phosphorylation and ubiquitination, and disruption of these posttranslational modifications can cause circadian dysfunction [5, 6]

Casein kinase 1 (CK1) is an evolutionally conserved core clock component [6]. The role of CK1 in the clock is complex. Mutations in CK1, first identified in hamsters [7, 8] and Drosophila [9, 10], can cause both period shortening and lengthening, while pharmacologic inhibition of CK1 consistently causes period lengthening in animals [11–13] and in plants [14]. Intriguingly, the kinase activity of CK1 on synthetic peptides is temperature-insensitive [12, 15]. This is consistent with a central role for CK1 in temperature compensation of the circadian period, the mechanism that maintains a stable circadian period despite changes in ambient temperature [16–18]. In mammals, CK1 delta and epsilon (CK1δ/ε) are essential kinases for the circadian clock [19]. The first identified mammalian circadian clock mutant, the *tau* hamster, harbors a semidominant mutation in CK1ε, and *tau* homozygous animals exhibit markedly shortened locomotor rhythms with a period of 20 h [7, 8]. In humans, mutations in CK1δ also cause Familial Advanced Sleep Phase (FASP) syndrome and shorten the circadian period [20, 21].

The biochemical function of CK1δ/ε in the clock arises from its stoichiometric tight interaction with and phosphorylation of PER proteins [4, 8, 22, 23]. One major consequence of PER2 phosphorylation by CK1δ/ε is the regulation of PER2 stability via the phosphoswitch [11, 17, 18, 24–27]. The PER2 phosphoswitch requires at least two phosphorylation domains. Cell-based and biochemical studies have established that Ser478 of PER2 protein is phosphorylated by CK1δ/ε, and Ser478 phosphorylation creates a phosphodegron that recruits an E3 ubiquitin ligase, beta-transducin repeat-containing homologue protein (β-TrCP) [11, 28]. β-TrCP is a substrate recognition subunit of the Skp1-Cullin-F-box (SCF) complex that catalyzes formation of a polyubiquitin chain on a substrate targeted for proteasomal degradation. The activity of CK1δ/ε on the S478 phosphodegron is controlled by a second PER2 phosphorylation domain called the FASP region. In this region, CK1δ/ε phosphorylates Ser659, followed by rapid phosphorylation of additional downstream serines in the motif pSxxSxxS. Cell based assays have demonstrated that this multi-phosphorylation of the FASP domain stabilizes PER2 by inhibiting phosphorylation of the S478 phosphodegron [17, 18, 24, 26]. Indeed, the FASP region is so named because a mutation of human PER2 Ser662 (corresponding to Ser659 in mouse) causes FASP syndrome [25, 29]. In the phosphoswitch model, the short-period phenotype common to *tau* and FASP mutations has been attributed to enhanced phosphorylation of the β-TrCP site leading to more rapid PER2 degradation. To date, however, there is no *in vivo* evidence that the β-TrCP site plays a role in determining the circadian period. To test this, we generated two knock-in mouse lines carrying Ser-to-Ala mutation at position 478 of PER2 protein. We find that PER2-Ser478Ala protein is stabilized in the mouse liver, and the mutation caused lengthening of the circadian period of mouse behavioral rhythms. These data provide the first genetic evidence for the importance of the β-TrCP site in the time-keeping mechanism of the circadian clock *in vivo.*

## Result

### The PER2-S478A mutation lengthens the period of behavioral rhythms

The phosphoswitch model holds that PER2 stability is bidirectionally regulated by two key phosphorylation sites, the β-TrCP site and the FASP site (Figure 1A). The model also predicts that a mutation of PER2 at the β-TrCP site should result in a more stable protein and a longer circadian period length *in vivo.* To test this hypothesis, we used CRISPR-Cas9 to generate a mouse with PER2 Ser478Ala (PER2-S478A)(Figure 1A). PER2-S478A homozygous mice were morphologically normal and fertile and were born in the expected Mendelian ratio. The mutant mice showed robust wheel-running activity rhythms under a 12 h light: 12 h dark (LD) cycle, and no apparent abnormality was observed in the light entrainment (Figure 1B). On the other hand, when transferred to constant dark (DD) condition, the mutant mice showed ~1 h longer period of the behavioral rhythms (24.58 ± 0.08 h) than wild-type (23.67 ± 0.06 h) (Figure 1C). This is the first *in vivo* demonstration that the Ser478 phosphodegron of PER2 determines the circadian period.

**Figure 1.**
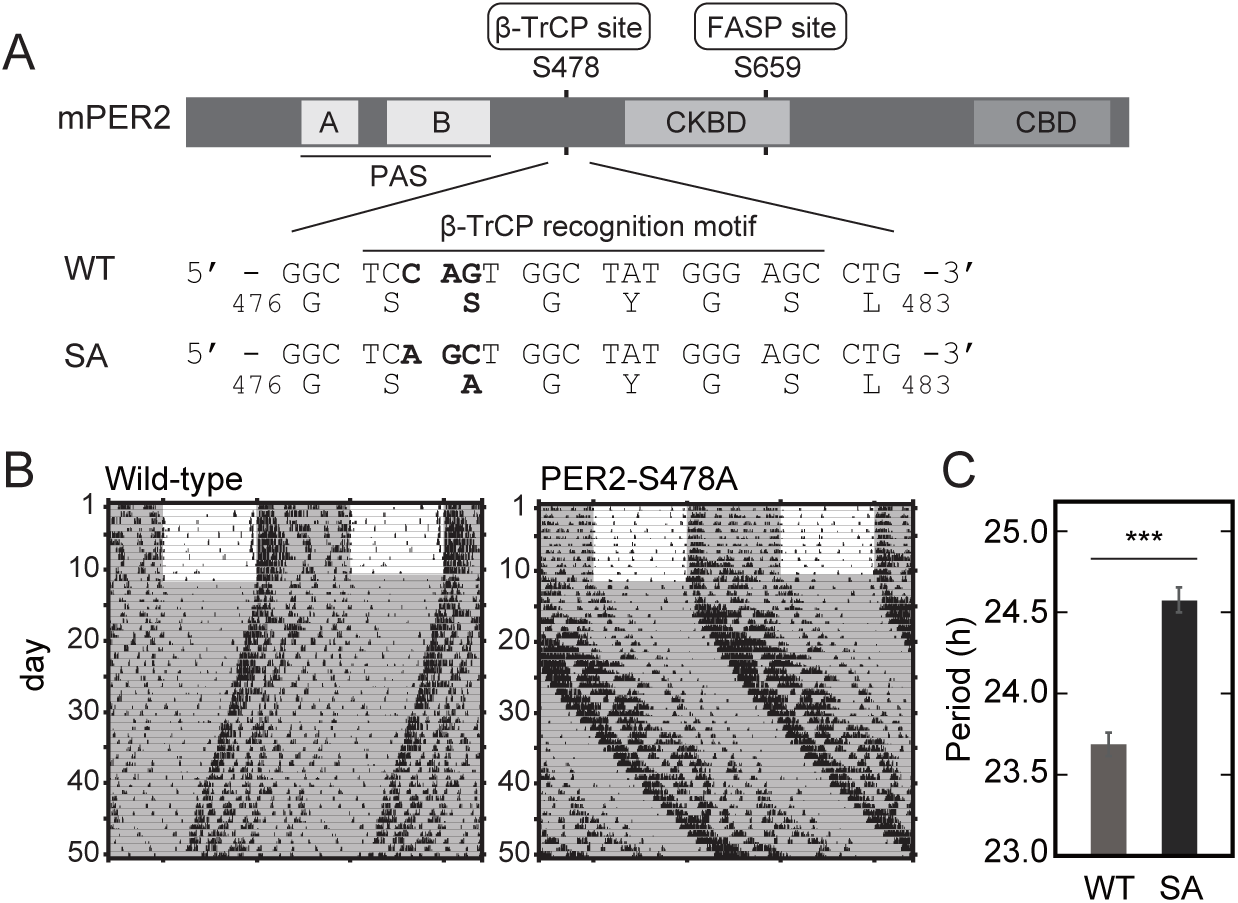
The PER2 Ser478Ala mutation lengthens circadian rhythms. (A) The domain structure of PER2 protein is depicted (upper). The genome sequences in wild-type (WT) and PER2-S478A (SA) knock-in mouse and corresponding amino acid sequences are shown (lower). PAS, Per-Arnt-Sim domain; CKBD, CK1-binding domain; CBD, CRY-binding domain. (B) Wheel-running activities of representative PER2-S478A homozygous mice and littermate wild-type mice are shown. Mice were entrained to LD for two weeks or longer and then exposed to DD. (C) The circadian period of the activity rhythms under the DD condition was determined via a chi-square periodogram procedure based on locomotor activity in days 11–24 after the start of DD condition. Mean values ± SEM obtained from ten wild-type (WT) mice and twelve homozygous PER2-S478A (SA) mice are given. *** *p* = 4.2 x E-8, two-sided Student’s *t*-test.

### Excessive accumulation of PER2-S478A mutant protein in the mouse liver

When over-expressed in cultured cells, PER2-S478A mutant was more stable than wild-type PER2 protein [17]. Here we examined the abundance of endogenous *Per2* mRNA and PER2 protein in the livers of PER2-S478A mice. Livers were harvested at six time points during the second day in DD condition. *Per2* transcripts showed the expected circadian oscillation in both wild-type and PER2-S478A mice, with no significant difference observed between the genotypes (Figure 2A). In contrast, peak levels of PER2 protein in both the nucleus and cytoplasm were ~2-fold higher at CT22 in PER2-S478A mice (Figure 2B and 2C). Thus, the S478A mutation stabilizes PER2 protein *in vivo* similar to what was seen in *in vitro* over-expression assays [17]. It should be noted that, although the peak levels of PER2-S478A protein were higher, PER2-S478A protein was still almost undetectable at CT10 and CT14, suggesting that phosphorylation of Ser478 is required for temporal degradation of PER2, and that other degradation mechanism(s) may be contributing as well [30].

**Figure 2.**
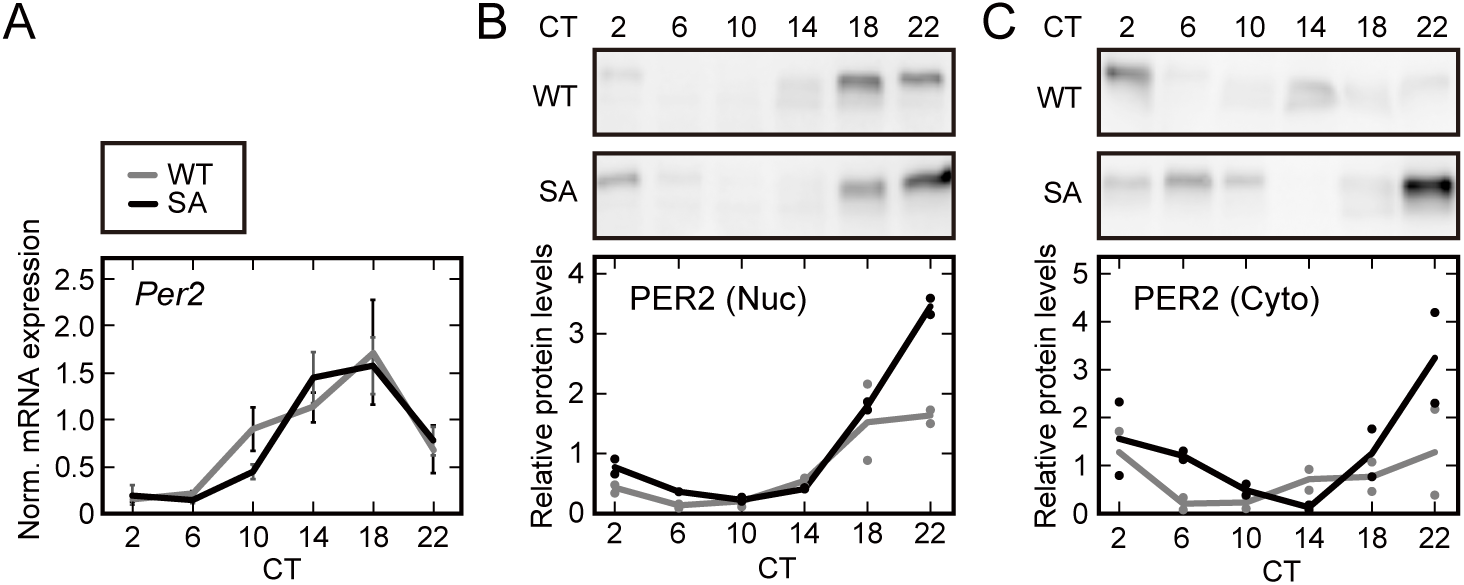
PER2 expression levels are post-transcriptionally upregulated in PER2-S478A mouse liver. (A) Temporal expression of *Per2* mRNA normalized by *Rps29.* Livers were harvested at 4-h intervals followed by real-time RT-qPCR. Mean values ± SEM obtained from three animals of each genotype are given. (B, C) Temporal expression of PER2 proteins in liver. Liver nuclear extracts (B) and cytoplasmic lysates (C) were prepared at 4-h intervals following SDS-PAGE and immunoblotting. Data are represented as dots for individual experiments and as lines for means.

### The PER2-S478A mutation stabilizes CRY proteins

PER2 has been proposed to be rate-limiting for *in vivo* formation of a stable multi-protein complex containing PER1, CRY1/2 and CK1δ/ε [2, 4] that assembles in the cytoplasm and then translocates to the nucleus. To assess the consequence of stabilization of PER2-S478A protein on these core circadian proteins, we examined the expression of a series of clock proteins in the livers of wild-type and mutant mice. At the time of peak PER2 abundance (CT22, Figure 2B, C), the cytosolic protein levels of PER1 and CRY2 were also increased (Figure 3A). Lagging by several hours, the nuclear abundance of PER1, CRY1 and CRY2 was also increased in the livers of PER2-S478A mice (Figure 3B). Thus, stabilized PER2 increases the abundance of the full circadian repressor complex.

**Figure 3.**
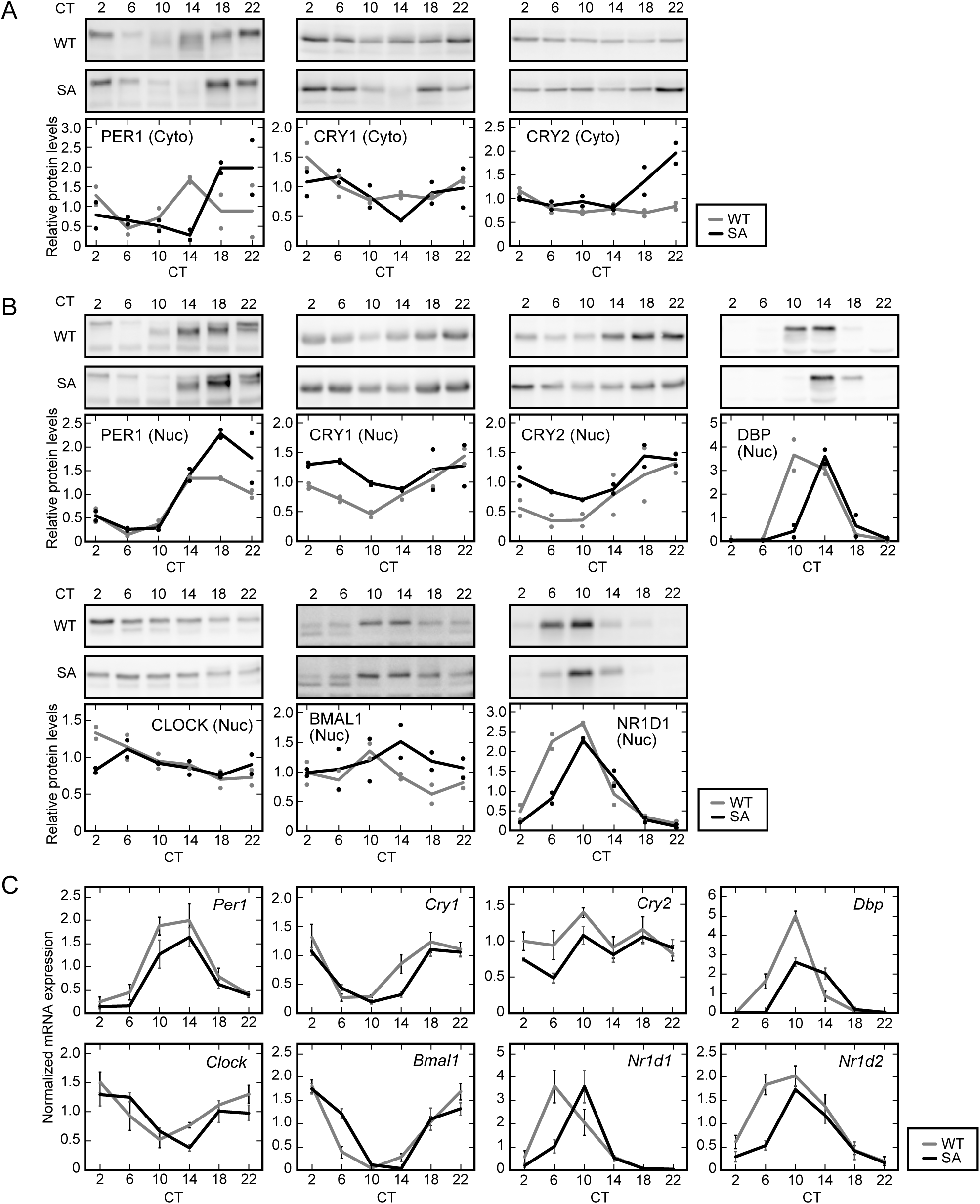
The PER2-S478A mutation alters the expression profiles of clock genes and proteins in the liver. (A, B) Temporal expression of clock proteins in mouse liver. Liver cytoplasmic lysates (A) and nuclear extracts (B) were prepared at 4-h intervals and analyzed by SDS-PAGE and immunoblotting with indicated antibodies. Data are represented as dots for individual experiments and as lines for means. (C) Temporal expression of clock genes. Livers were harvested at 4-h intervals followed by real-time RT-qPCR for the indicated mRNAs. Mean values ± SEM obtained from three animals of each genotype are given.

Several studies proposed that PER2 stabilizes CRY proteins by interfering with the interaction between CRYs and FBXL3, an E3 ubiquitin ligase that directs degradation of CRYs [31–33]. Up-regulation of CRY protein levels in the PER2-S478A mutant liver (Figure 3A, B) suggests that CRYs are stabilized in response to elevation of PER2 protein levels. To test this, we measured the half-lives of CRY1::LUC and CRY2::LUC fusion proteins in HEK 293T cells by monitoring the bioluminescence decay after treatment of cycloheximide (CHX) inhibiting new protein translation. When PER2-S478A was co-expressed, CRY1::LUC and CRY2::LUC were stabilized in a dose-dependent manner and to the same extent as wild-type PER2 protein (Figure S1). Thus, stabilization of PER2 appears to stabilize CRY proteins in both cultured cells and in mouse liver.

PER abundance has been proposed to be the rate-limiting step in the assembly and nuclear entry of the repressor complex [2, 4, 34]. Notably, the nuclear protein levels of CRY1 and CRY2 were still high at CT10 (Figure 3B) when a large proportion of PER2-S478A protein was degraded (Figure 2B). This suggests that CRY proteins can repress CLOCK-BMAL1 transcriptional activity even in the absence of PER proteins, consistent with a previous study [35]. In fact, the mRNA expression profiles of typical E-box-regulated genes, such as albumin D-site binding protein *(Dbp), Nr1d1* (also known as *Rev-erbA)* and *Nr1d2* (also known as *Rev-erbB*), were phase-delayed in the PER2-S478A mutant liver (Figure 3C). The delayed expression profiles were also observed in RRE-regulated genes *Bmal1, Clock,* and *Cry1* (Figure 3C). These profiles are consistent with the longer behavioral rhythms of the PER2-S478A mice. Concomitantly, nuclear expression profiles of DBP and NR1D1 proteins were delayed in the mutant mouse liver (Figure 3B). Overall, our data indicate that the PER2-S478A mutation results in accumulation of circadian repressor complex in the nucleus, leading to delayed de-repression of the CLOCK-BMAL1 activator.

### The S478A mutation of PER2 compromises temperature compensation

The phosphoswitch model arose from the observation that PER2::LUC degradation occurred in three phases [17]: an initial rapid decay (the first phase), a plateau-like slow decay (the second phase), and finally, a more rapid terminal decay (the third phase). This was visualized by continuous monitoring of bioluminescence signals from PER2::LUC knock-in MEFs. To investigate how the S478A mutation affects the three-phase degradation *in vivo,* we in parallel introduced the S478A mutation into the PER2::LUC allele (PER2-S478A::LUC) encoding a PER2 fusion protein with firefly luciferase [36]. In the analysis of the wheel running activity rhythms, PER2-S478A::LUC mice exhibited a significantly longer circadian period than PER2::LUC control mice (Figure 4A, B), similar to that seen in the PER2-S478A mice. We established mouse embryonic fibroblasts (MEFs) from both PER2-S478A::LUC and control PER2::LUC mice.

**Figure 4.**
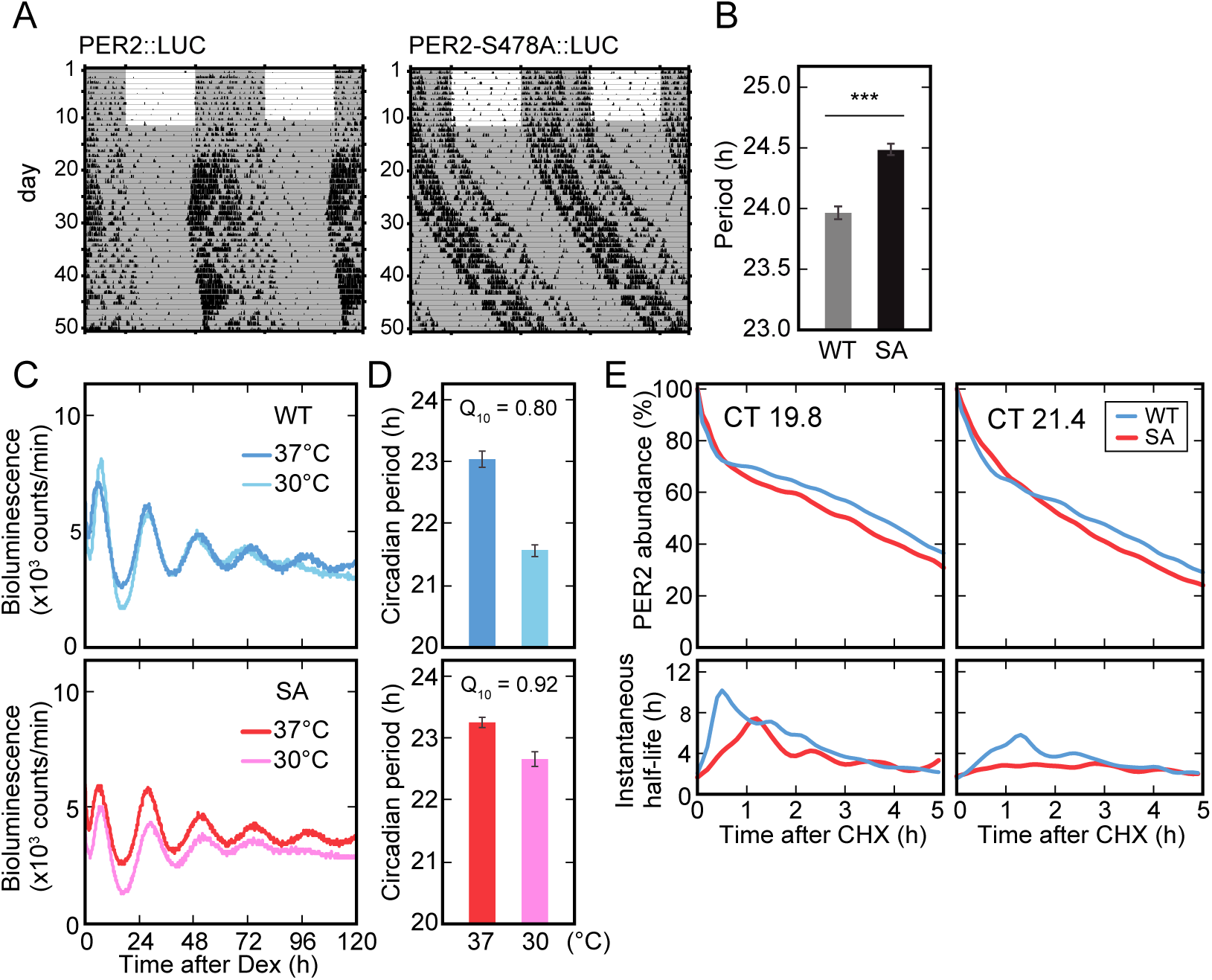
The mutation of PER2 at Ser478 perturbs the phosphoswitch. (A) Wheel-running activities of representative PER2::LUC and PER2-S478A::LUC homozygous mice are shown. Mice were entrained to LD for two weeks or longer and then exposed to DD. (B) The circadian period of the activity rhythms under the DD condition was determined via a chi-square periodogram procedure based on locomotor activity in days 11–24 after the start of DD condition. Mean values ± SEM obtained from ten PER2::LUC mice (WT) and ten PER2-S478A::LUC (SA) mice are given. *** *p* = 5.6 x E-7, two-sided Student’s *t*-test. (C) Representative recordings of bioluminescence from PER2::LUC or PER2-S478A::LUC MEFs are shown. MEFs were synchronized with dexamethasone and the bioluminescence was then continually measured in the Lumicycle. (D) The circadian period of cellular rhythms was calculated. Mean values ± SEM obtained from three PER2::LUC MEFs (WT) and three PER2-S478A::LUC (SA) MEFs are given. (E) Bioluminescence from PER2::LUC (WT) and PER2-S478A::LUC (SA) MEFs are shown. MEFs were synchronized with dexamethasone and the bioluminescence was continually measured in the Lumicycle. CHX was added at CT 19.8 (left) or at CT 21.4 (right). By fitting the exponential curve to the small segment of the decay curves, the instantaneous half-life is calculated (see Materials and methods for details).

We then tested if disruption of the phosphoswitch by the S478A mutation affects the temperature compensation, which is a key feature of the circadian clock [37]. While the rate of most enzymatic reactions doubles or triples with every 10°C increase in temperature *(i.e.,* temperature coefficient, Q_10_ = 2-3), circadian rhythms are relatively stable in the face of environmental temperature alterations, and in fact are often over-compensated, with a Q_10_ < 1.0. Indeed, in the PER2::LUC MEFs, as the temperature decreased from 37°C to 30°C the clock sped up ~1.5 hours, giving a Q_10_ = 0.80 (Figure 4C, D). However, in the PER2-S478A::LUC MEFs, the clock only sped up ~0.5 hours, for a Q_10_ of 0.92 (Figure 4C, D). Thus, temperature compensation was compromised by the S478A mutation, underscoring the importance of this site in the phosphoswitch-regulated temperature compensation.

Finally, we examined whether mutation of the phosphodegron can disrupt the previously observed three-phase decay of PER2::LUC. When protein translation was inhibited by addition of cycloheximide (CHX) at two distinct time points (CT19.8 and CT21.4) in the PER2 accumulation phase, we again observed three-phase degradation of endogenous PER2::LUC (Figure 4E). The duration of the second phase was dependent on the time when CHX was added, as previously reported [17]. At both time points, the three-phase degradation of PER2 protein was markedly attenuated by the S478A mutation (Figure 4E), resembling what was seen when CK1 was inhibited [17].

Collectively, these data demonstrate that the phosphodegron in PER2 plays a key role in the phosphoswitch, temperature compensation and circadian period *in vivo*.

## Discussion

Robust genetic data indicate that CK1δ/ε controls the speed of the circadian clock oscillation, for which multiple potential molecular mechanisms have been proposed. These include nuclear-cytoplasmic shuttling of PER proteins, phosphorylation of PER-associated proteins in the repression complex, phosphorylation of CLOCK to decrease its DNA binding, and a phosphoswitch to regulate PER protein stability [4, 17, 38]. We previously demonstrated that phosphorylation of the FASP region inhibited phosphorylation of the β-TrCP phosphodegron, but it has not been experimentally addressed whether the β-TrCP degron actually regulates the circadian period. In the present study, we find that phosphorylation of the β-TrCP site controlling PER2 stability is an essential step in determining the period length *in vivo.* Mice with the PER2-S478A mutation showed significantly longer behavioral rhythms than control mice (Figure 1). The period lengthening effect (by ~1 h) is similar to those observed in mice injected with a CK1 inhibitor [13] and in transgenic mice carrying human PER2 S662D, a phospho-mimic mutant at the FASP site [25]. These data show that the β-TrCP site is important for the regulation of behavioral rhythm by the phosphoswitch. Unlike what was observed in hPER2-S662D transgenic mice [25], *Per2* mRNA levels were unaltered in the PER2-S478A mutant. The peak levels of PER2 protein were increased in cytosol and nucleus (Figure 2), consistent with disruption of the Ser478 phosphodegron. Thus, this study provides a strong genetic evidence that phosphorylation-regulated degradation of PER2 is indeed a key regulator of the clock speed.

CK1δ and CK1ε may not be the only kinases regulating PER2 stability. For example, a chemical screening identified CK1α as a regulator of PER protein stability [39]. This effect of CK1α may be indirect because CK1α binds to PER proteins with a much lower affinity than CK1δ/ε does [38–40]. In *Drosophila,* CK1α enhances PER degradation elicited by DBT (the CK1δ/ε homolog in *Drosophila*) [41]. CK2 phosphorylates PER2 at Ser53 and promotes PER2 proteasomal degradation in both CK1δ/ε-dependent and-independent pathways [42]. PKCα [43] and GRK2 [44] regulate the nuclear entry of PER2 and might affect the stability of PER2. In this study, we showed that PER2-S478A mutation attenuated CK1δ/ε-mediated degradation of PER2 protein and lengthened the circadian period. Loss of function of CK1 in *Neurospora* also lengthened the circadian period [45]. The clock genes are not orthologous across species in eukaryotes, but architectures of the molecular clockworks are analogous. FREQUENCY (FRQ), a negative component of the circadian clock in *Neurospora,* is analogous to PER. CK1 phosphorylates FRQ and decreases its protein stability as well [46]. In contrast, recent studies proposed that the binding affinity of CK1 for FRQ, instead of FRQ stability, might be a critical factor for the period determination through phosphorylation of other clock proteins [47, 48]. Our data establish a role for the phosphodegron in regulating circadian rhythms but do not rule out an alternative model in which CK1-mediated phosphorylation of other substrates also plays a role for regulation of clock timing.

In this study, we propose a model (Figure 5), in which PER2 Ser478 phosphorylation governs decay of PER/CRY-containing repressor complex that determines restart timing of the E-box-dependent transcriptional activation. The peak level of PER2 protein is regulated by the balance between the competing phosphorylations of stabilizing (FASP) and destabilizing (β-TrCP) sites. The PER2-S478A mutation disrupts this balance and results in excessive PER2 accumulation (Figure 2) that interferes with the degradation of PER1 and CRY1/2 proteins (Figure 3). The mature repressive complex enters the nucleus to inhibit CLOCK-BMAL1. The nuclear PER proteins are then degraded, presumably in a β-TrCP site-independent fashion. The remaining CRY proteins stay on CLOCK-BMAL1 activator complex to keep repressing the transcriptional activity in the absence of PERs [35, 50, 51], also contributing to the lengthened period phenotype (Figure 1).

**Figure 5.**
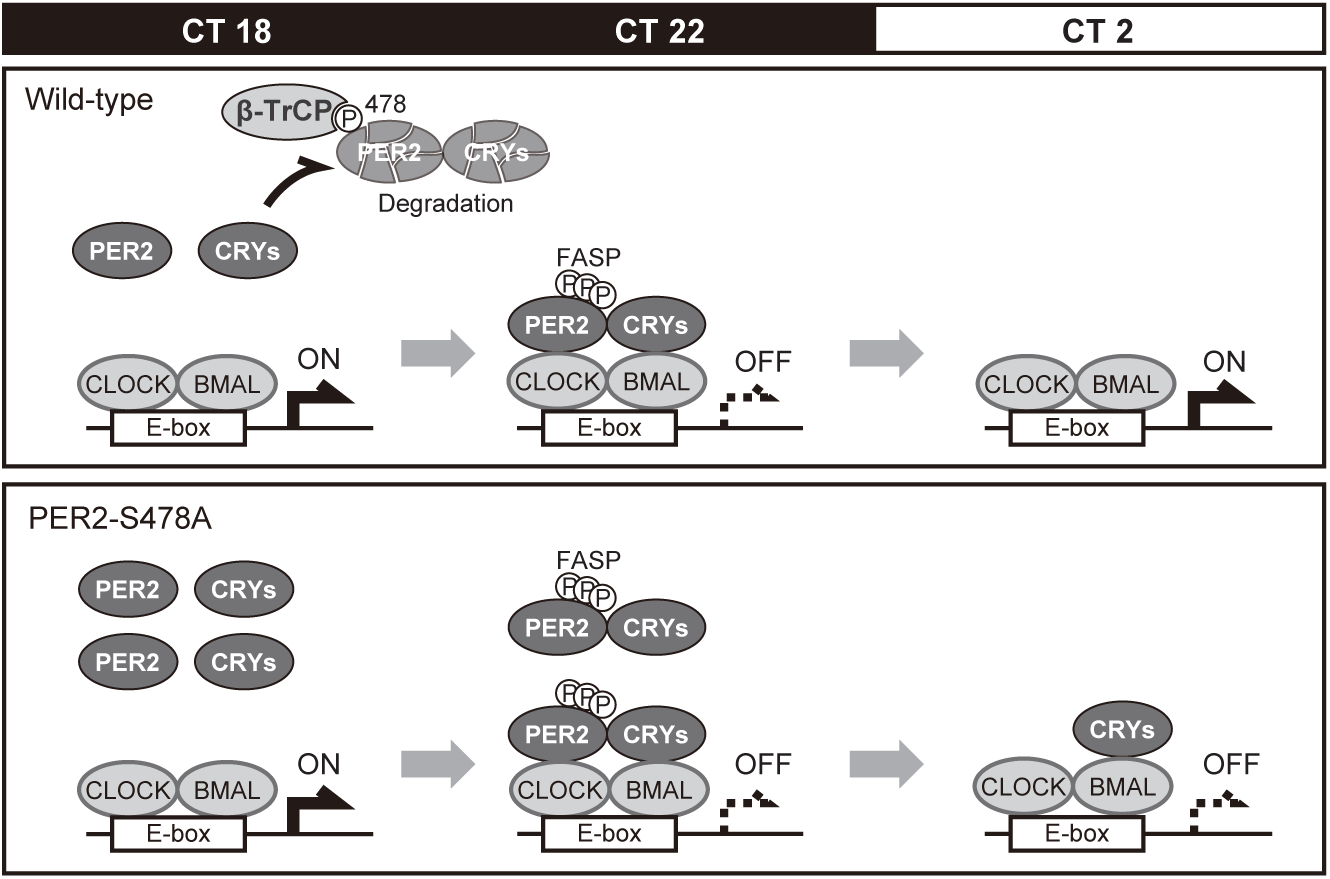
PER2-S478A mutation lengthens the circadian period. PER2 and CRY proteins accumulate during night and disappear before CT2 in wild-type. In contrast, PER2 and CRY proteins are stabilized and excessively accumulate around CT 22 in PER2-S478A mutants. Thereafter, PER2 proteins are degraded by CT2 whereas CRY proteins keep repressing the CLOCK-BMAL complexes.

Temperature compensation (or over-compensation) of the circadian period has been described for more than 60 years [37]. Mathematical models predicted that temperature-dependent opposing reactions contribute to the temperature-compensated clockwork [49], although the processes which virtually act in this mechanism have not been understood. We previously demonstrated that the PER2 phosphoswitch may act as the compensating process in mammalian clockwork [17]. The phosphoswitch leads to two pools of PER2, one less stable due to degron phosphorylation, and the other more stable due to FASP phosphorylation inhibiting degron phosphorylation. Therefore, three-phase degradation can be observed following CHX treatment in PER2::LUC MEFs [17]. Here, the three-phase degradation was compromised in PER2-S478A::LUC MEFs, consistent with this model. The mutant MEFs showed also cellular rhythms with a longer period at lower temperature. These results are consistent with the idea that the β-TrCP site is more susceptible to phosphorylation at lower temperature to make the circadian period compensated for temperature changes [17].

Although mutation of the Ser478 phosphodegron increased PER2 stability, PER2 was still degraded between CT22 and CT2 (Figure 2). Furthermore, the three-phase degradation was indeed perturbed in the PER2-S478A mutant but was not completely abolished (Figure 4E, left panels). These data suggest alternative pathway(s) for PER2 degradation. One possible pathway is via an additional β-TrCP site. A potential phosphodegron motif TpSGCSpS is conserved in PER1 (aa121-126) and PER2 (aa92-97), whereas the primary β-TrCP degron SpSGYGpS (aa477-482 in PER2) is not present in PER1 [52]. In PER2, the phosphorylation of the primary degron plays a dominant role for β-TrCP-mediated degradation, but a role of the second degron has been reported when the primary degron is mutated [28]. In PER2-S478A mutant, this additional degron may therefore contribute to the alternative degradation pathway. Second, it was recently reported that oncoprotein MDM2 targets PER2 for proteasomal degradation, independent of the phosphorylation status of PER2 protein [30]. Third, it is possible that β-TrCP recruited to phosphorylated PER1 might contribute to PER2 degradation in the repressor complex. Future studies on detailed mechanisms underlying the PER2 stability control will help to understand the temperature compensation of the circadian clock.

## Acknowledgements

We thank Dale Cowley of TransViragen/UNC Chapel Hill Animal Models Core for design and production of the Ser478Ala mutant mice, and Dr. Atsu Aiba in the University of Tokyo for their help with mouse embryo freezing and the embryo transfer. This work was partially supported by Singapore’s NMRC/CIRG/1465/2017 to D.V., by Grants-in-Aid for Specially Promoted Research to Y.F. (17H06096), and for Scientific Research (B) to H.Y. (25440041) from MEXT of Japan, and by the PRIME from Japan Agency for Medical Research and Development to H.Y. (17937210).

## Author contributions statement

S.M., R.N., H.Y., Y.F., and D.V. designed the study. S.M. performed the almost all experiments. R.N. performed the three-phase degradation assay and J.K.K analyzed it. H.Y., Y.F., and D.V. supervised the whole project. S.M. and H.Y. wrote the first draft of the manuscript. Y.F. and D.V. edited the manuscript. All authors reviewed and approved the final manuscript.

## Materials and methods

### Mice

Mice used in this study (C57BL/6 background) were handled in accordance with the Guidelines for the Care and Use of Laboratory Animals at The University of Tokyo. Mice were housed in cages with free access to commercial chow (CLEA Japan) and tap water. The animals were maintained in a light-tight chamber at a constant temperature (23°C ± 1°C) and humidity (65% ± 10%).

### Generation of PER2-S478A mice and PER2-S478A::LUC mice

Four Cas9 guide RNAs targeting the mouse *Per2* Ser478 codon were selected for activity testing. Guide RNAs were cloned into a T7 promoter vector followed by *in vitro* transcription (HiScribe T7 High Yield RNA Synthesis Kit, New England BioLabs) and spin column purification (RNeasy, Qiagen). Functional testing was performed by incubating Cas9 protein and guide RNA with PCR-amplified target site followed by gel electrophoresis to detect target site cleavage. The guide RNA selected for genome editing in embryos was Per2-sg72T (protospacer sequence 5’-GCGGCTCCAGTGGCTAT-3’). Donor oligonucleotide Per2-S478A-T was used as a donor for point mutation insertion in the *Per2* gene (5’-CTTCTACACCTTTCTATGTGCTTTCCCCCCAGCCTGTCCCCCACAGCGGCTCAGCTGGCTATGGGAGCCTGGGCAGTAACGGATCCCACGAACACCTCATGAGCCAGA −3’). SpCas9 protein was expressed and purified by the UNC Chapel Hill Protein Expression and Purification Core Facility.

Embryos were produced by in vitro fertilization with C57BL/6J females (Jackson Laboratory) as egg donors and *Per2^luc/luc^;CK1ε^+/+^;CK1δ^flox/flox^* males as sperm donors. Single-cell fertilized embryos were microinjected with 400 nM Cas9 protein, 50 ng/ul guide RNA and 50 ng/ul donor oligonucleotide in microinjection buffer (5 mM Tris-HCl pH 7.5, 0.1 mM EDTA). Injected embryos were implanted in recipient pseudopregnant females and resulting pups were screened by PCR and sequencing for the presence of the *Per2^S478A^* allele. Founders harboring the *Per2^S478A^* allele were mated to wild-type C57BL/6J females for germline transmission of the targeted allele and to determine if the targeted allele was in cis with the luciferase knock-in. Note that because the *Per2^luc^* allele was produced by gene targeting in ES cells from strain 129, the *Per2* locus in the embryos was heterozygous with 129-derived and C57BL/6J-derived alleles. PCR amplicon sequencing documented polymorphisms in the vicinity of the Ser478 codon that could be used to distinguish the 129 and C57BL/6J alleles. Founders with the S478A mutation in the C57BL/6J allele were used to generate animals with the S478A mutation without the luciferase. Founders with the S478A allele inserted in cis with luciferase in the 129 allele were used to establish the combined S478A-Luc strain.

For the establishment of the Ser478Ala mutant lines, biopsy DNA was genotyped by PCR amplification of a 1,042 bp region encompassing the Ser478 codon with primers Per2-ScF2 (5’-CGGGTCTCTCTGTGCACTCTTG −3’) and Per2-ScR2 (5’-GATGCTTCCTTCTGTCCTCCA −3’). The PCR product was sequenced with primer Per2-SqR2 (5’-GTGACTTTTGGTTTGAATCTTGC −3’). For genotyping, DNA was extracted from the tail and analyzed by the polymerase chain reaction with the primers Per2-ScF2, Per2-ScFwt (5’-CACAGCGGCTCCAGTGG-3’), Per2-ScRmut (5’-AGGCTCCCATAGCCAGCTG-3’) and Per2-ScR3 (5’-GCTTCTCAGGGAGAGGAACAG-3’).

### Behavioral experiments

Six-to ten-week-old male mice were individually housed in cages equipped with running wheels and were entrained to the 12-h /12-h LD cycle. After more than two weeks, wheel revolutions were recorded under the DD condition for 24 days or longer. Their spontaneous locomotor activities were recorded as the number of wheel revolutions in 5-min bins and were analyzed with ClockLab software (Actimetrics). The circadian period of the activity rhythms under the DD condition was determined via a chi-square periodogram procedure from animals that showed rhythmicities with P < 0.001, based on locomotor activity in days 11–24 after the start of DD condition.

### Cell culture and real-time monitoring of rhythmic gene expression

MEFs from PER2::LUC mice and PER2-S478A::LUC mice were maintained at 37°C under 5% CO2, 95% air in Dulbecco’s modified Eagle’s medium (SIGMA) supplemented with 100 units/ml penicillin, 100 μg/ml streptomycin, and 10% fetal bovine serum. We carried out real-time monitoring of luciferase expression as previously described [53] with minor modifications. Briefly, cells were treated with 0.1 μM (final) dexamethasone for 30 min, and then the media were replaced by recording media [phenol-red free Dulbecco’s modified Eagle’s medium (SIGMA) supplemented with 10% fetal bovine serum, 3.5 g/l glucose, 25 U/ml penicillin, 25 μg/ml streptomycin, 0.1 mM luciferin, and 10 mM HEPES-NaOH; pH 7.0]. The bioluminescence signals were continuously recorded for 5-10 days at 37°C in air with Dish Type Luminescencer, LumiCycle (Actimetrics).

### RNA preparation

Tissues from 7–8-week-old male mice were collected every 4 h from 38 h after the beginning of the DD condition (projected circadian time (CT)). Livers with three biological replicates were lysed with TRIzol reagent (Invitrogen), and total RNA was prepared with the RNeasy Mini Kit (Qiagen) according to the manufacturer’s protocol.

### RT-qPCR

For quantification of gene expression, RNA was reverse-transcribed by Go Script Reverse Transcriptase (Promega) with both anchored (dT) 15 primers and random oligo primers. We subjected the cDNA to real-time PCR (StepOnePlus Real-Time PCR Systems, Applied Biosystems) using GoTaq Master Mix (Promega) with the gene specific primers (Table 1).

**Table 1.**
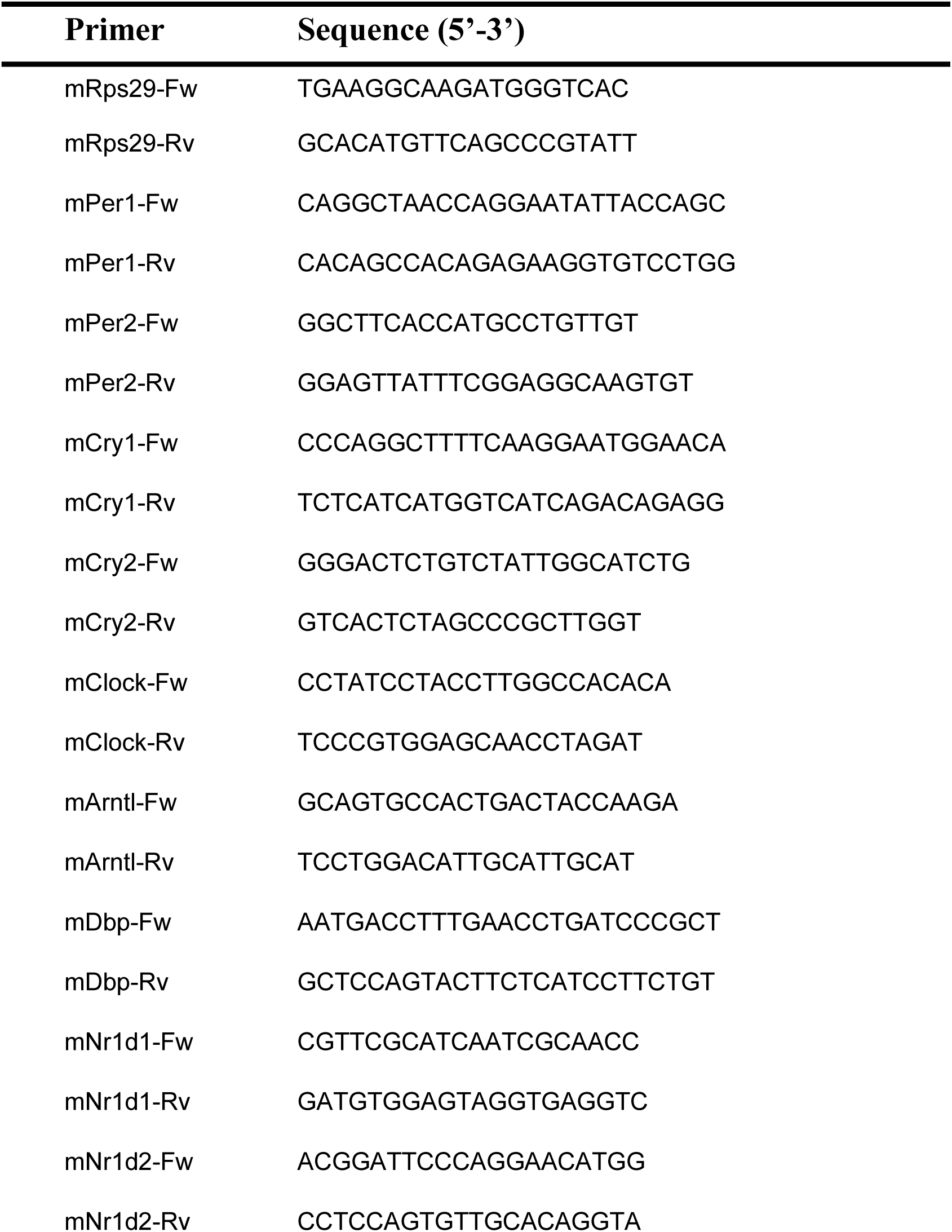
Primer sequences for RT-qPCR

### Preparation of nuclear and cytosolic fractions of mice liver

The nuclear proteins and cytosolic proteins were isolated as previously described [54]. Briefly, the mouse tissue (1 g wet weight) was washed with icecold PBS and homogenized on ice with 11 ml of ice-cold buffer A [10 mM HEPES-NaOH (pH 7.8), 10 mM KCl, 0.1 mM EDTA, 1 mM dithiothreitol (DTT), 1 mM phenylmethylsulfonyl fluoride (PMSF), 4 μg/ml aprotinin, 4 μg/ml leupeptin, 50 mM NaF, and 1 mM Na_3_VO_4_]. The homogenate was centrifuged (5 min each, 700 xg), and the supernatant was used as “cytosolic fraction”. the resultant precipitate was rinsed and centrifuged (5 min each, 700 xg). The precipitate was resuspended in 2 ml of ice-cold buffer C [20 mM HEPES-NaOH (pH 7.8), 400 mM NaCl, 1 mM EDTA, 5 mM MgCl_2_, 2% (v/v) glycerol, 1 mM DTT, 1 mM PMSF, 4 μg/ml aprotinin, 4 μg/ml leupeptin, 50 mM NaF, and 1 mM Na_3_VO_4_].

After being gently mixed at 4°C for 1 h, the suspension was centrifuged (30 min each, 21,600 x g) twice, and the final supernatant was used as “nuclear fraction”.

### Antibodies and immunoblot analysis

Liver nuclear and cytosolic proteins separated by SDS-PAGE were transferred to polyvinylidene difluoride membrane (Millipore). The blots were blocked in a blocking solution (1%[w/v] skim milk in TBS [50 mM Tris-HCl, 140 mM NaCl, 1 mM MgCl_2_ (pH 7.4)]) for 1 h at 37°C and then incubated overnight at 4°C with a primary antibody in the blocking solution. The signals were visualized by an enhanced chemiluminescence detection system (PerkinElmer Life Science). The blot membrane was subjected to densitometric scanning and the band intensities were quantified using ImageQuant TL (GE Healthcare). The primary antibodies used in this study are as follows; anti-CLOCK [54] (CLSP3, Medical & Biological Laboratories, D333-3), anti-ARNTL [54] (B1BH2, Medical & Biological Laboratories, D335-3), anti-PER1 (Medical & Biological Laboratories, PM091), anti-PER2 (Medical & Biological Laboratories, PM083), anti-CRY1 (Medical & Biological Laboratories, PM081), anti-CRY2 (Medical & Biological Laboratories, PM082), anti-DBP (Medical & Biological Laboratories, PM079) and anti-NR1D1 (Medical & Biological Laboratories, PM092).

### Degradation assay

Degradation assays were performed as previously described [55] with minor modifications. Plasmids driving expression of CRY1::LUC or CRY2::LUC were co-transfected with myc-PER2 or myc-PER2-S478A plasmids in HEK 293T cells. After 48 h, the transfected cells were treated with 100 μg/ml cycloheximide (Nakalai tesque) before monitoring the bioluminescence in LumiCEC (Churitsu), or were harvested for immunoblotting. Degradation assay in cultured MEFs was performed in essentially the same way except that DNA was not transfected. To calculate half-lives of CRY1::LUC and CRY2::LUC, each bioluminescence data was fitted by two-term exponential model;

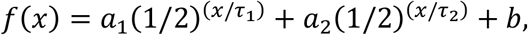

where x represent time, a_i_, b and τ_i_ are constant value such as initial quantities, baseline and half-lives, respectively. These values were computed by linear least square fittings. The fact that the data were fitted well by two-term exponential model indicates CRY1::LUC and CRY2::LUC can exist at least in two states with different stability. The half-life of unstable form (τ_1_) was likely to be independent of PER2 abundance, whereas that of stable form (τ_2_) was significantly increased in a PER2 dependent manner. The accumulation of CRY proteins in PER2-S478A mice liver can be explained by stabilization of the stable form of CRY proteins, thus, we described τ_2_ as the half-lives in Figure S1.

To calculate the instantaneous half-life of PER2, we divide the PER2 decay curve into segments of 1h with a sliding window, which is moved at increments of 0.1 h as done in Zhou et al [17]. Then, each segment of decay curve is fitted to the exponential decay curve.

---

**Supplemental Figure S1. PER2 stabilizes CRY1::LUC and CRY2::LUC**

(A, B) Half-lives of the bioluminescence derived from CRY1::LUC (A, upper panel) or CRY2::LUC (B, upper panel) after CHX treatment. The detailed method for calculation of half-lives is described in Materials and methods. Myc-PER2 (WT) or Myc-PER2-S478A (SA) were co-transfected with CRY1::LUC or CRY2::LUC. The abundance of PER2 proteins were analyzed by immunoblotting with anti-PER2 antibody (A, B, lower panel). Mean values ± SEM obtained from three biological replicates are given.

(C) Representative western blot for PER2, corresponding to (A, lower panel).

(D) Representative western blot for PER2, corresponding to (B, lower panel).

