## Supplementary figures and images for "Mutation of a PER2 phosphodegron perturbs the circadian phosphoswitch"

### Supplemental Figure 1

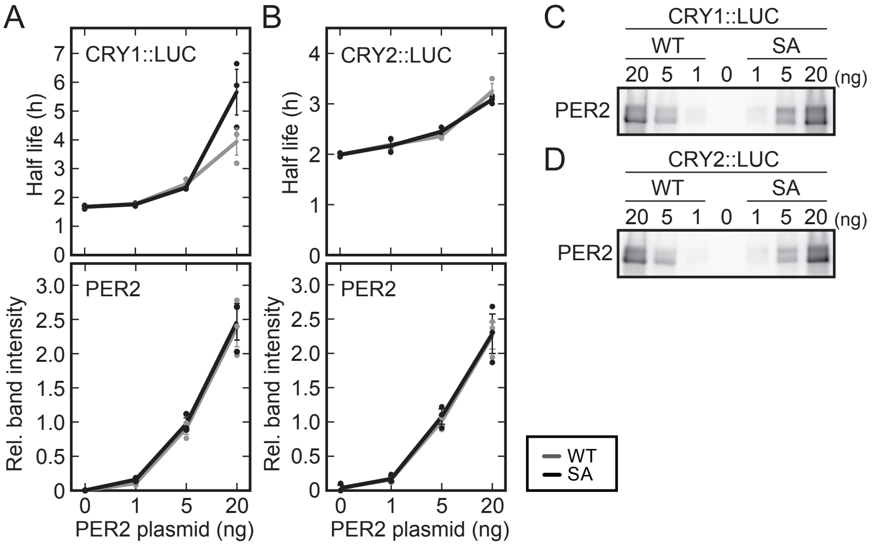
